## Supplemental Text 1 for "Trial-by-trial learning of successor representations in human behavior"

### Parameter Recovery

We wanted to verify that our model was capable of recovering distinct parameter regimes for parameters of interest. We verified parameter recovery through the following procedure. For each simulation, we chose a set of mean values for $\lambda$, $\gamma$, $\beta_A$, and $\beta_W$. We exhaustively simulated all combinations of $\lambda$ in [0.1, 0.3, 0.5, 0.7, 0.9], $\gamma$ in [0.5, 0.7, 0.9], $\beta_A$ in [-2, -1, 0], $\beta_W$ in [-2, -1, 0], treating each as the mean value of that parameter in our simulated dataset. For each of these 5 x 3 x 3 x 3 combinations of parameter values, we created a new set of 100 subjects. To create this set of subjects, we first sampled the best fit parameters for a single subject (from the experimental data), with replacement. Then, for that subject and for each of the four target parameters, we sampled the target parameter from a normal distribution centered at the target group mean with variance equal to the recovered group variance from our best fit model in the manuscript. We then generated synthetic reaction times, sampling log RTs from N($\mu$,$\sigma$), where $\mu$ is the predicted mean log RT from our simulation and $\sigma$ is the value fit for that individual subject. This process was repeated for all 100 subjects. Finally, we re-fit the model and all free parameters to this new simulated dataset. In figure S1, we plot the range of recovered group-level parameters across all simulations. We collapse across all parameter combinations: the recovered values of $\lambda$ for $\lambda = 0.1$ include all possible combinations of other parameters. We find that for all combinations of graphs and parameters, the recovered group-level parameters successfully captured the simulated means and were clearly distinguishable.

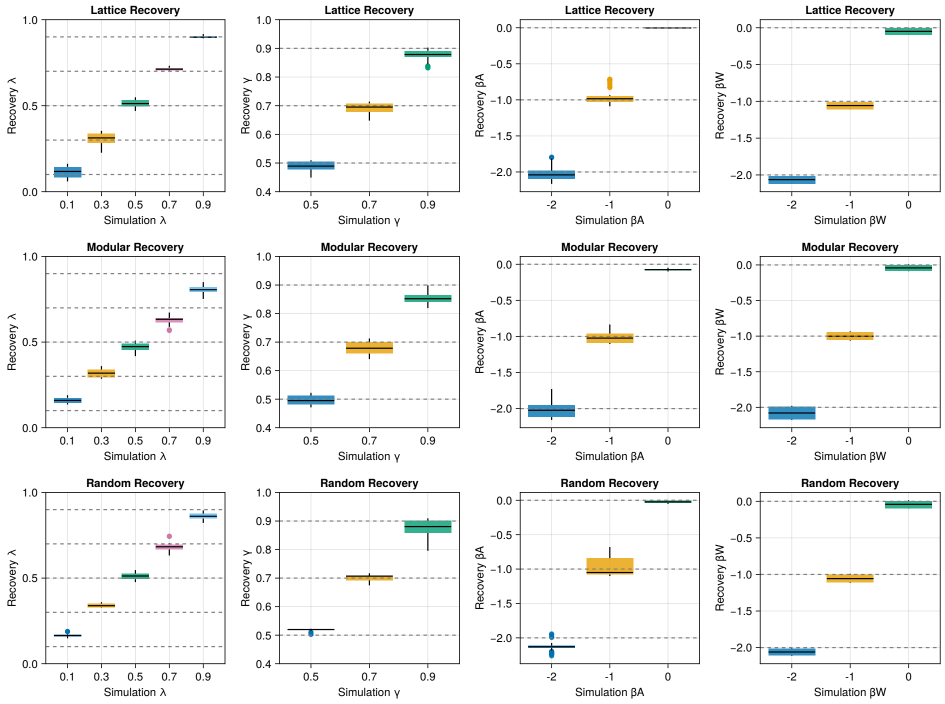


**Figure A. Parameter Recovery**. Boxplots indicate median, 25th, and 75th percentiles, and whiskers indicate 1.5 * interquartile range. Outlying points are plotted separately.

#### Subject Parameters

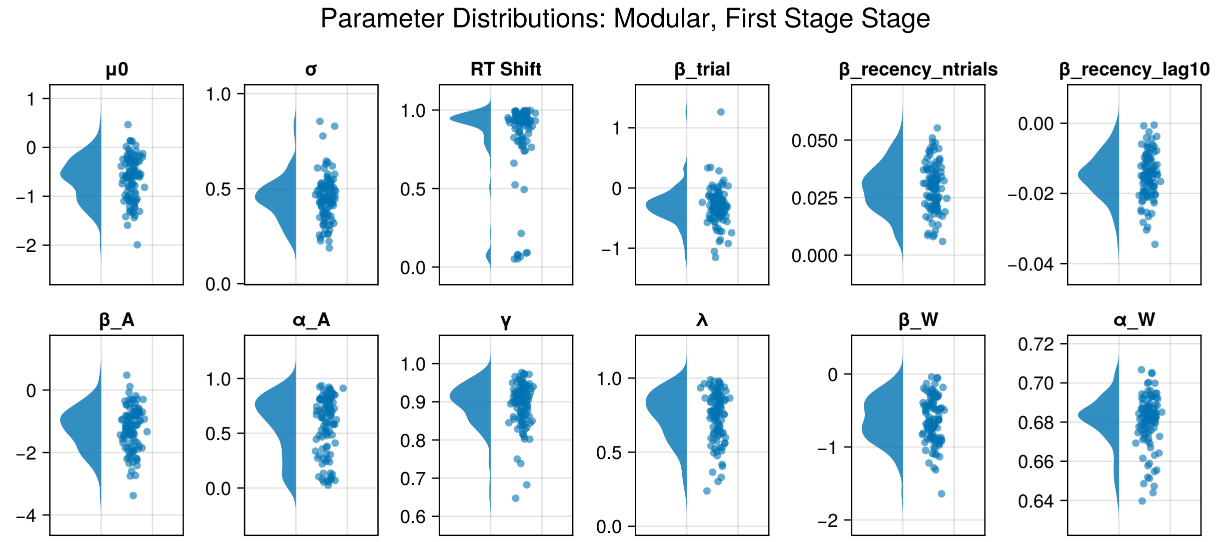


**Figure B.** Recovered parameters for First Stage Modular subjects. Each point is a single subject.

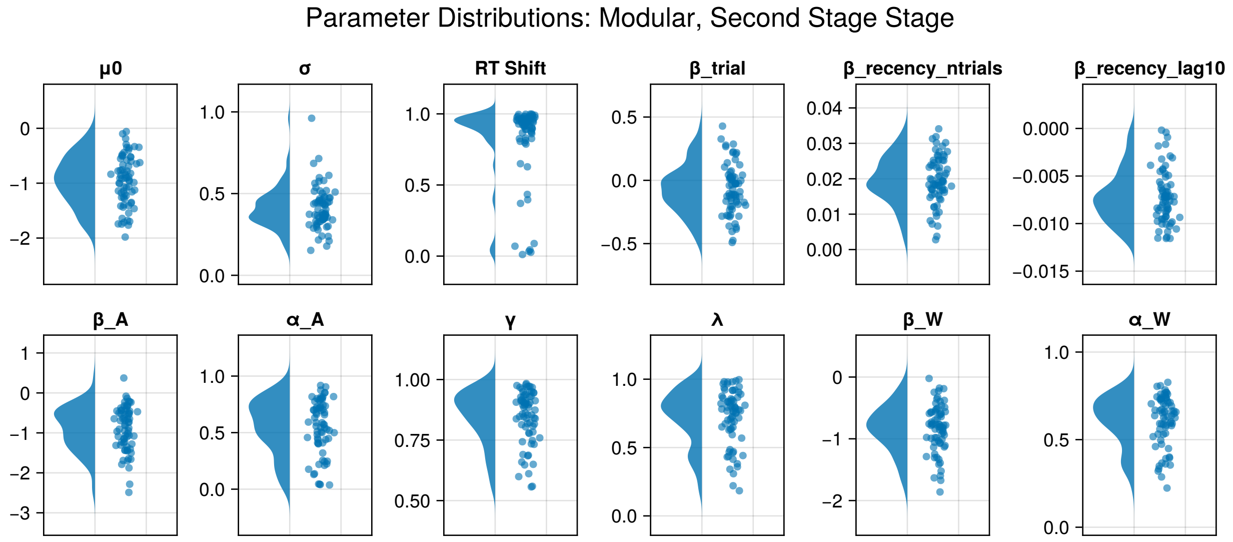


**Figure C.** Recovered parameters for Second Stage Modular subjects. Each point is a single subject.

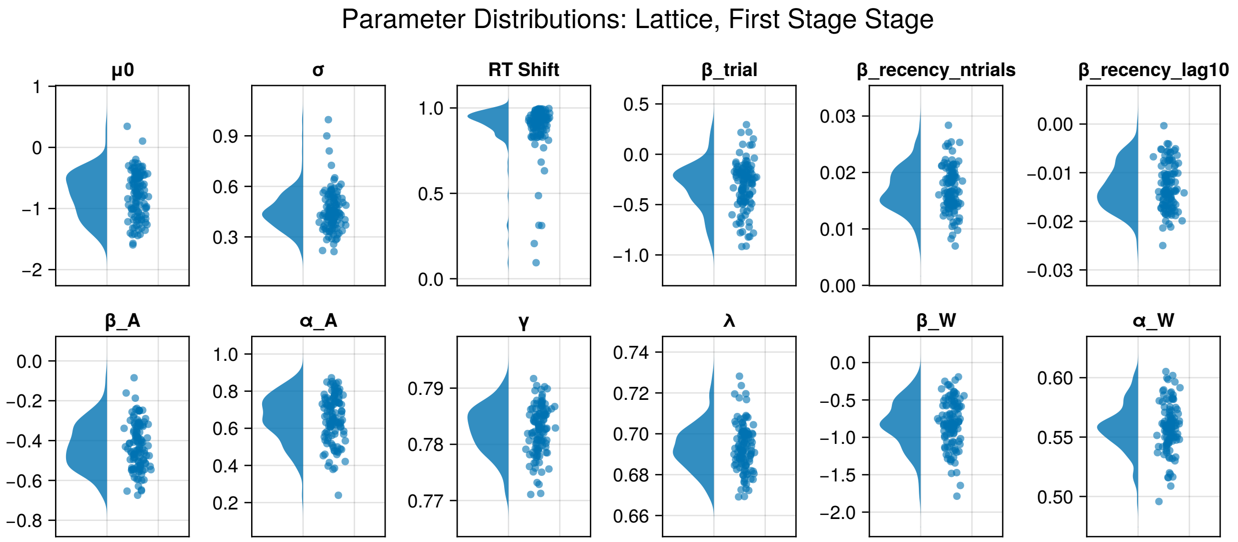


**Figure D.** Recovered parameters for First Stage Lattice subjects. Each point is a single subject.

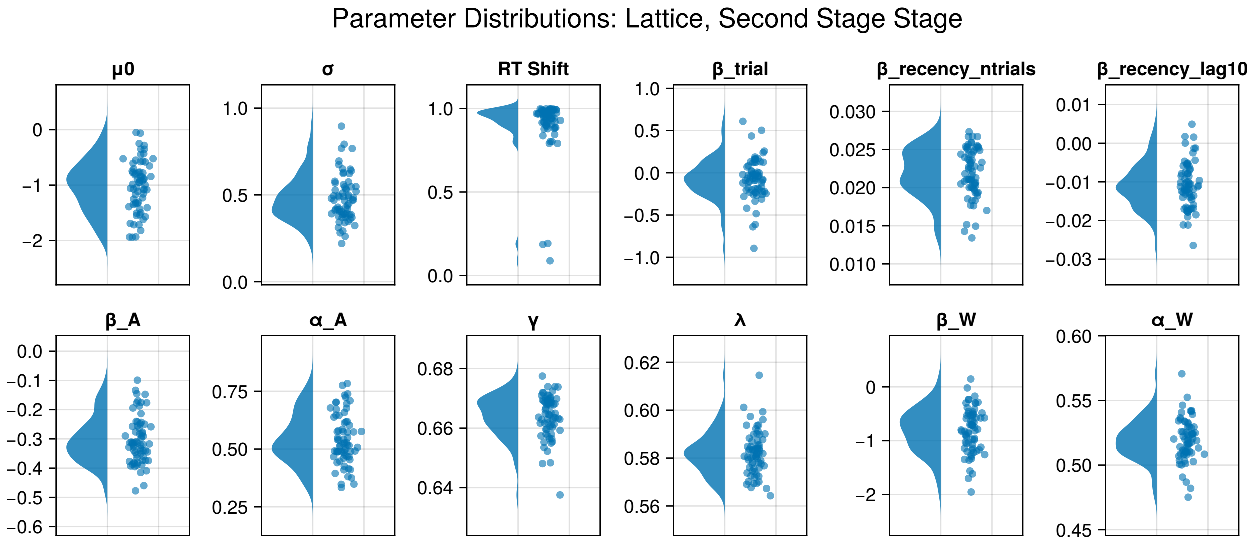


**Figure E.** Recovered parameters for Second Stage Lattice subjects. Each point is a single subject.

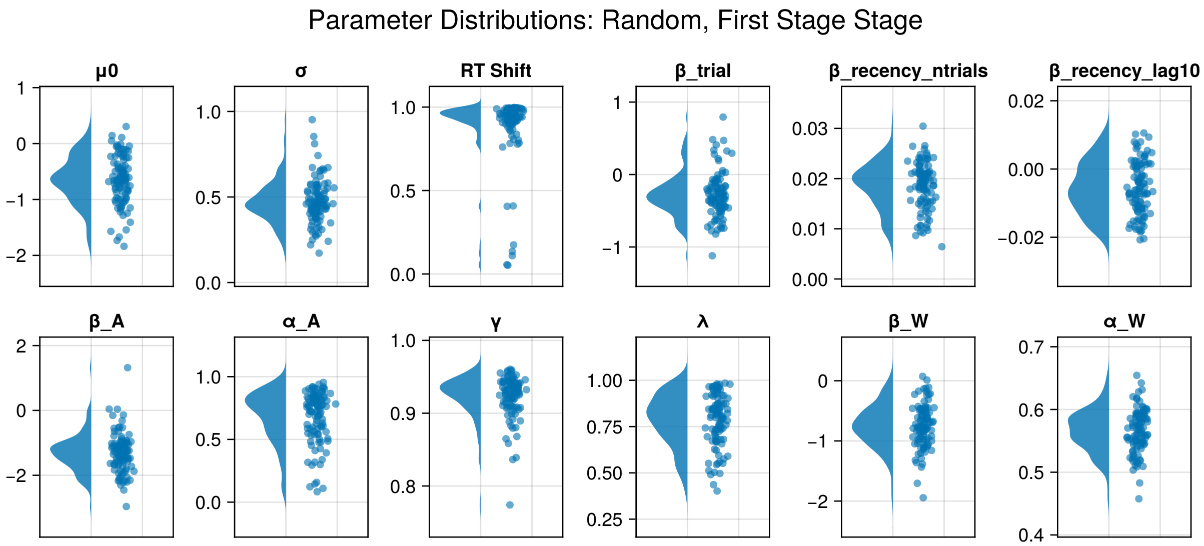


**Figure F.** Recovered parameters for First Stage Random subjects. Each point is a single subject.

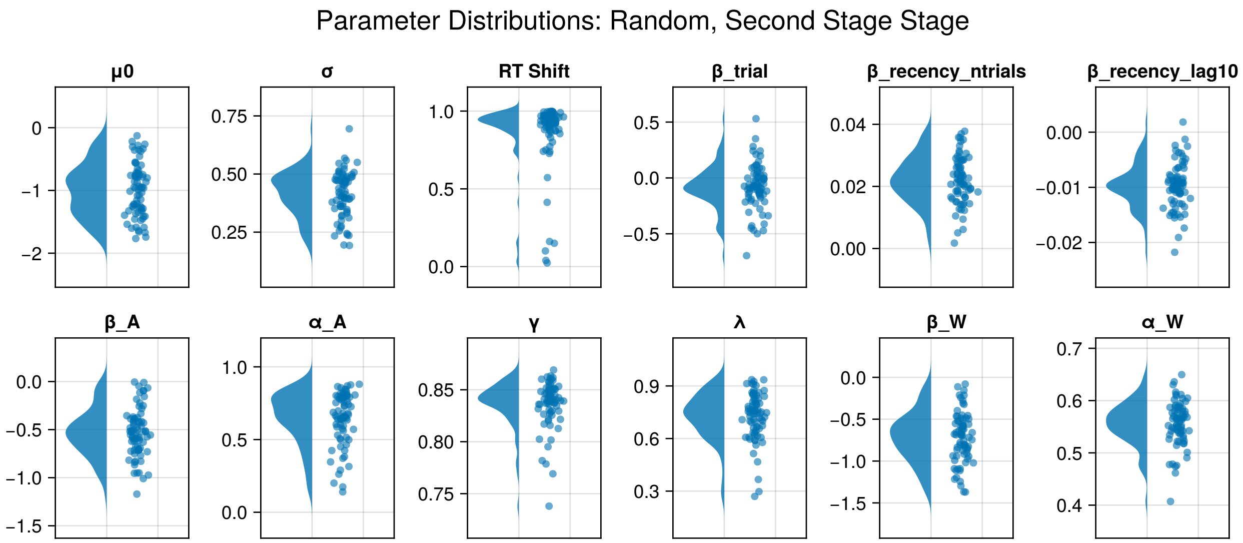


**Figure G.** Recovered parameters for Second Stage Random subjects. Each point is a single subject.

### Parameter Pair Plots

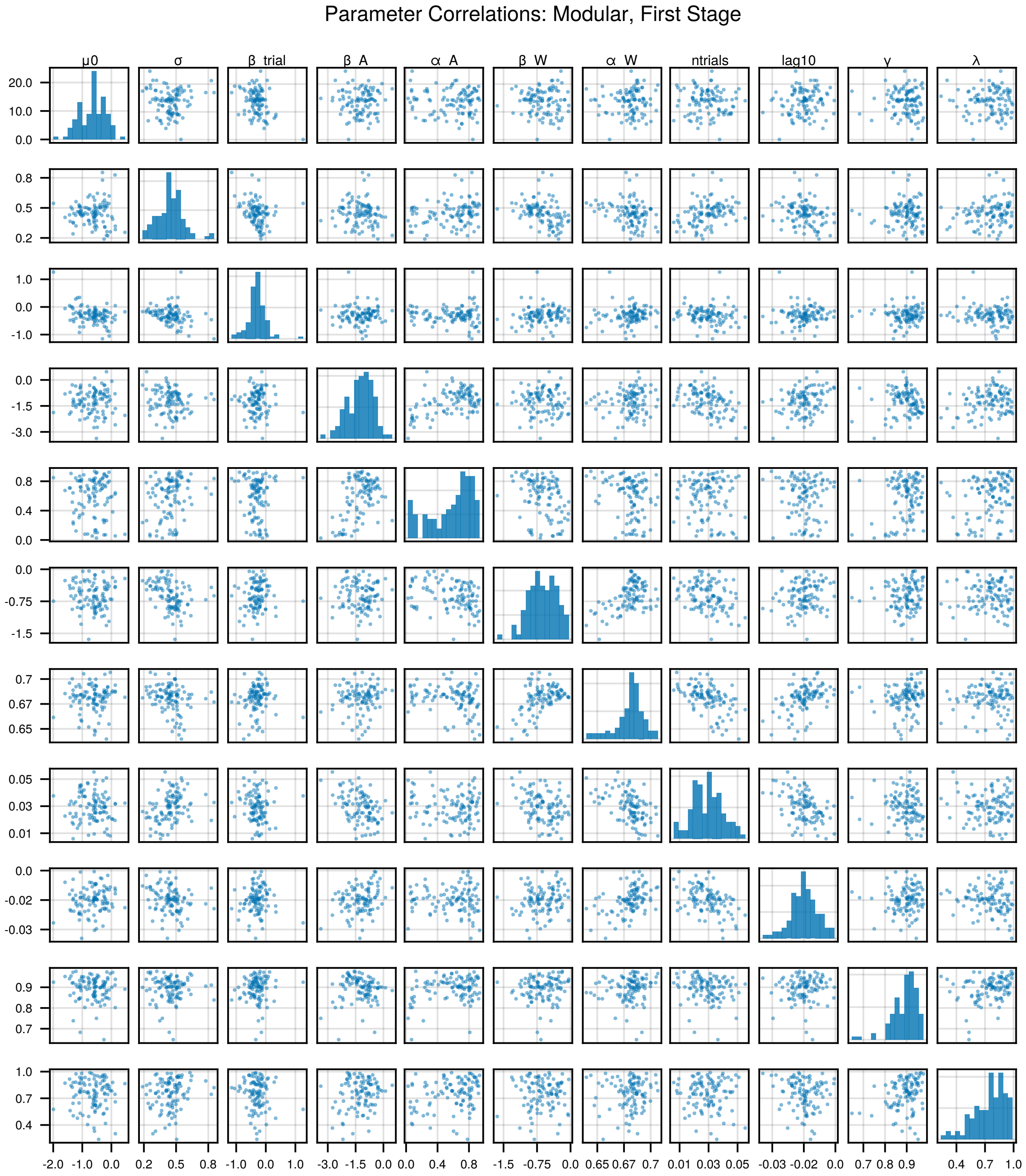


**Figure H.** Pair plots of all recovered parameters for First Stage Modular subjects.

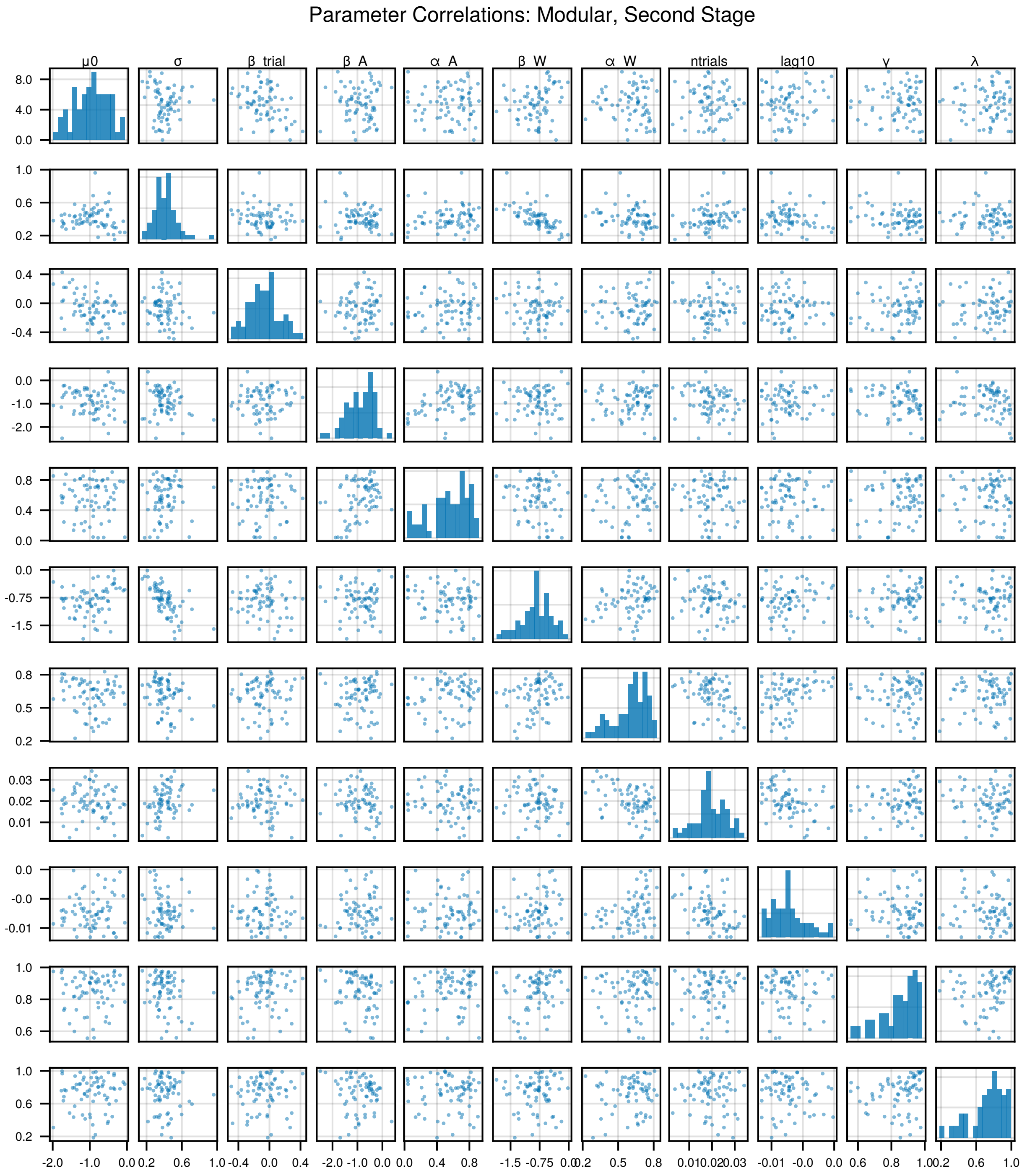


**Figure I.** Pair plots of all recovered parameters for Second Stage Modular subjects.

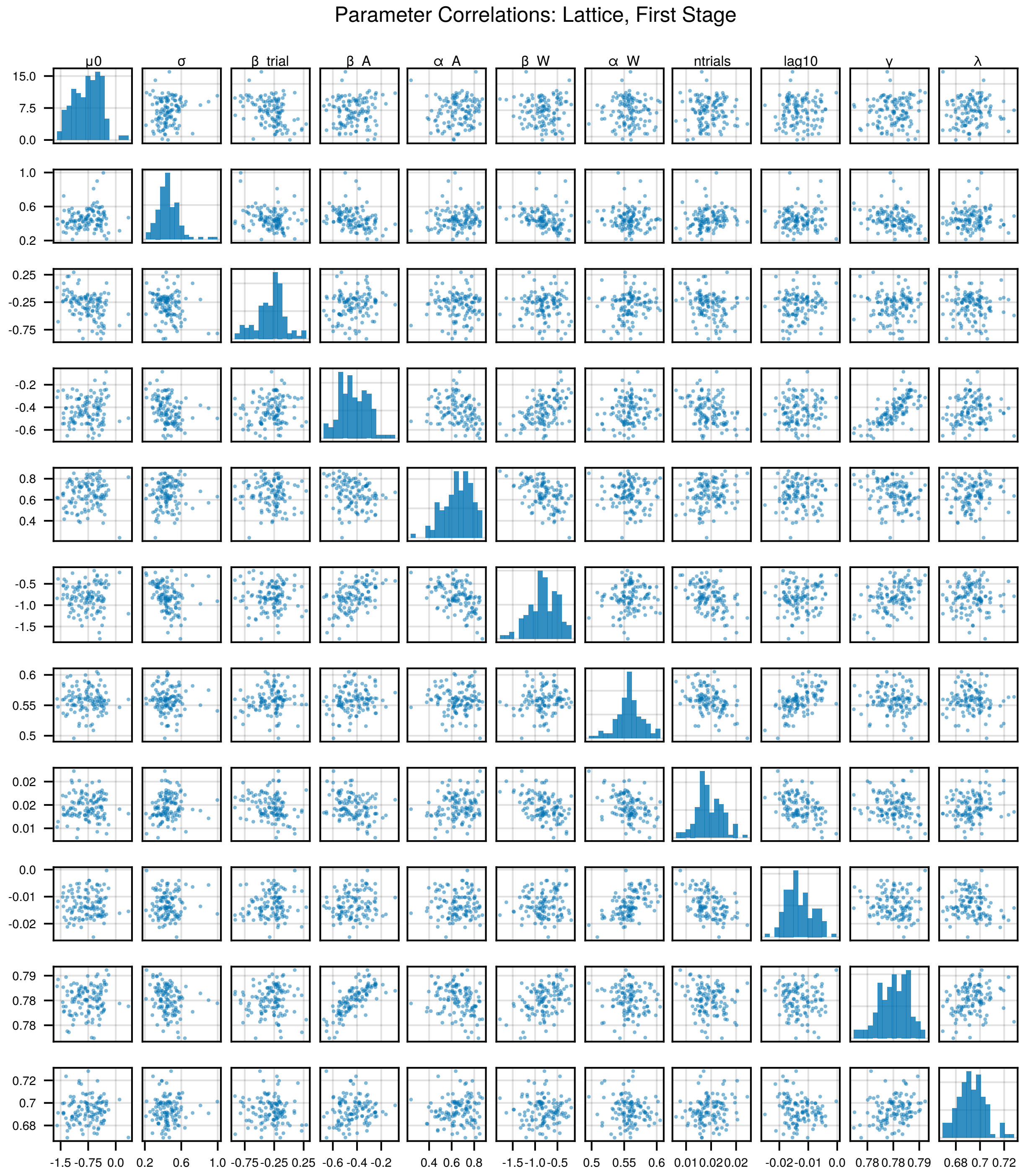


**Figure J.** Pair plots of all recovered parameters for First Stage Lattice subjects.

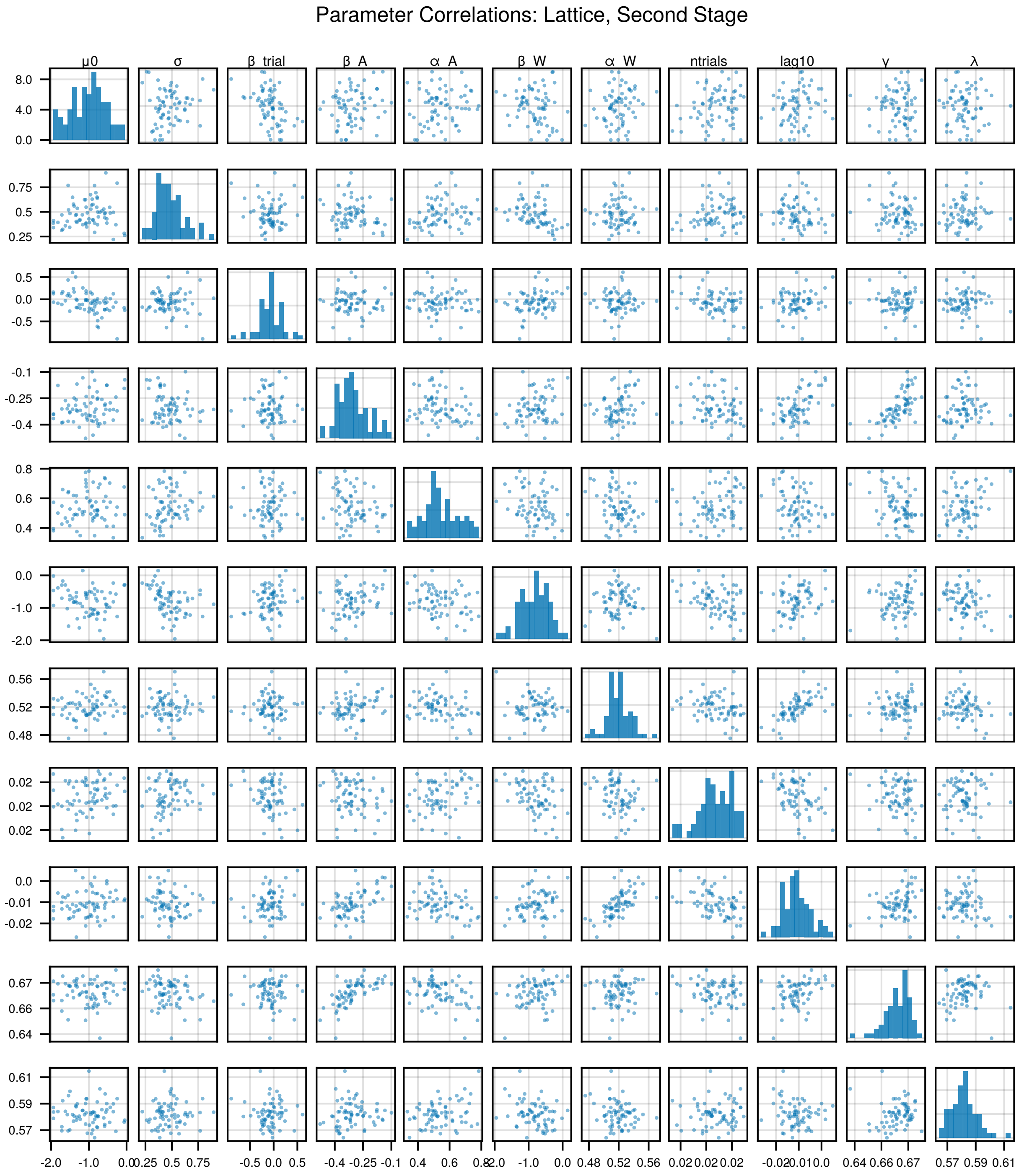


**Figure K.** Pair plots of all recovered parameters for Second Stage Lattice subjects.

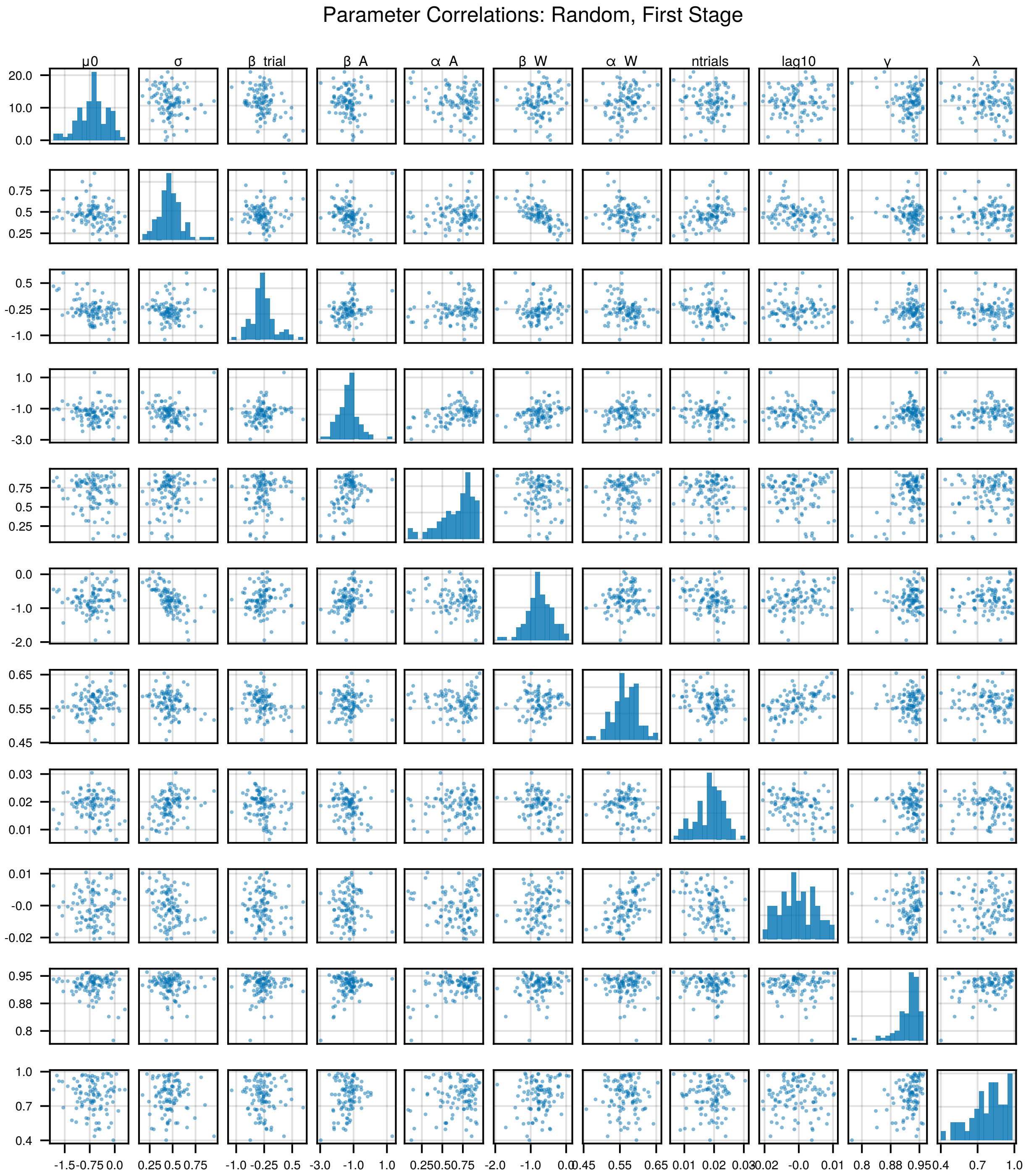


**Figure L.** Pair plots of all recovered parameters for First Stage Random subjects.

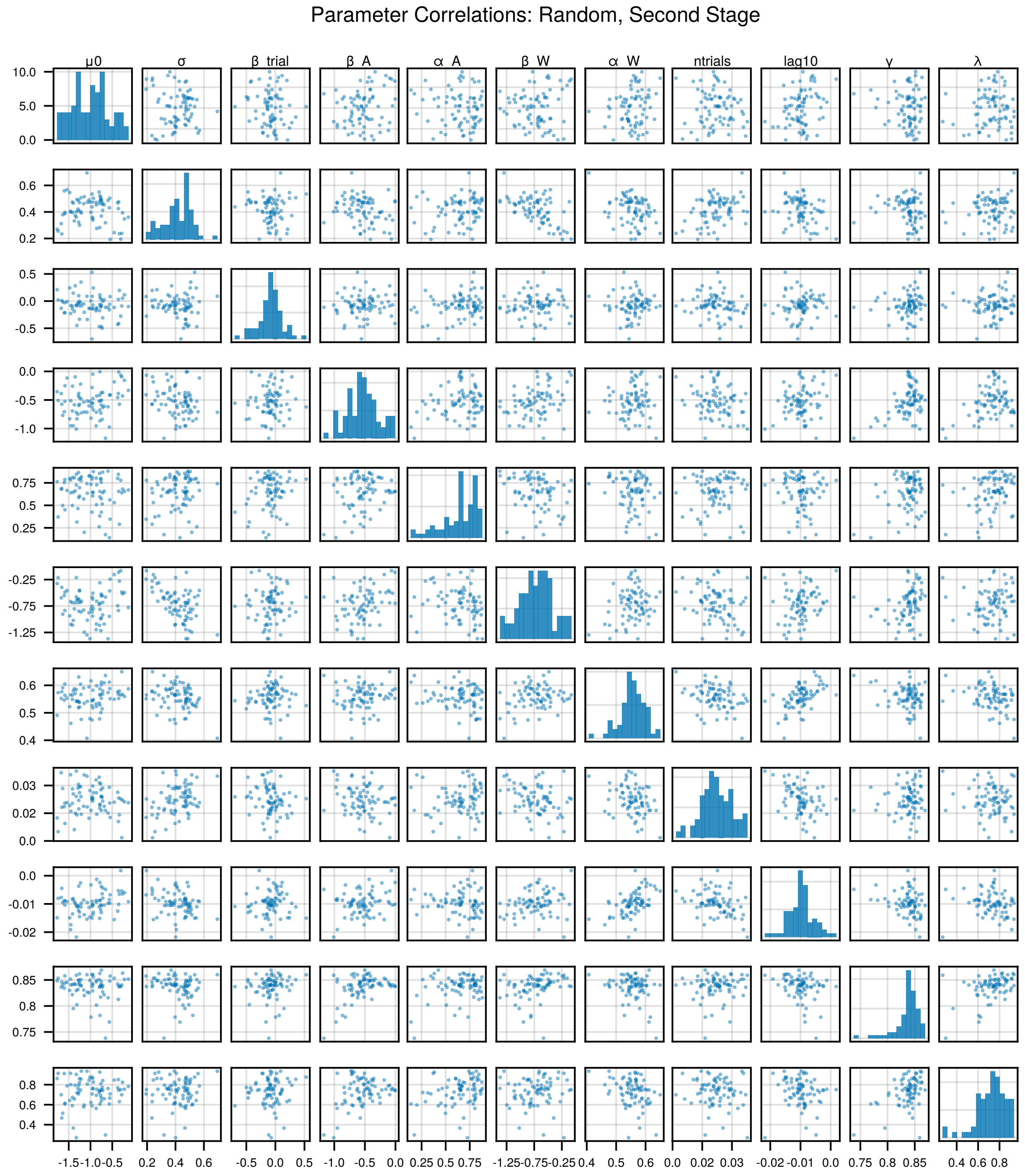


**Figure M.** Pair plots of all recovered parameters for Second Stage Random subjects.

### Model Distinguishability

To test that our models were distinguishable and able to capture unique predicted sources of variance, we simulated RTs using a TD0, TD1, or TD$\lambda$ model. For each simulated subject, we sampled best fit parameters of a single subject with replacement from a model fit to our experimental data, with either TD0, TD1, or TD$\lambda$, and used those parameters and the associated model to generate new RTs, sampling log RTs as in our parameter recovery. We then fit data from 100 such subjects, using either the TD0, TD1, or TD$\lambda$ model. We generated 20 unique datasets for each combination of simulation and recovery model. Plotted in figure S2 are the differences in fit versus the best fitting model for each simulation, measured via IAIC. We found that, for each combination of simulation and recovery, recovery with the appropriate model led to the lowest IAIC score for 19/20 or 20/20 simulations in all conditions.

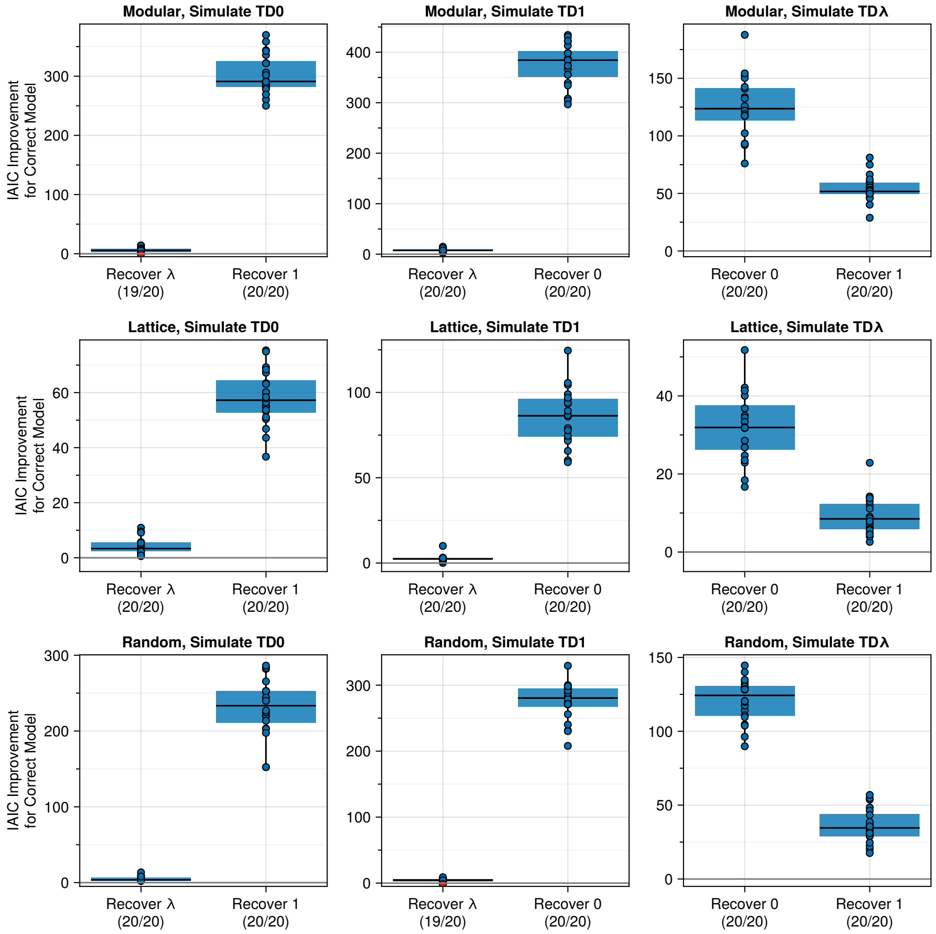


**Figure N. Model distinguishability.** Each point is the improvement in IAIC for recovery with the data simulation model versus an alternative model. Boxplots indicate median, 25th, and 75th percentiles, and whiskers indicate 1.5 * interquartile range.
